# In vivo imaging of swimming micromotors using hybrid high-frequency ultrasound and photoacoustic imaging

**DOI:** 10.1101/2020.06.15.148791

**Authors:** Azaam Aziz, Joost Holthof, Sandra Meyer, Oliver G. Schmidt, Mariana Medina-Sánchez

## Abstract

The fast evolution of medical micro- and nanorobots in the endeavor to perform non-invasive medical operations in living organisms boosted the use of diverse medical imaging techniques in the last years. Among those techniques, photoacoustic (PA) tomography has shown to be promising for the imaging of microrobots in deep-tissue (ex vivo and in vivo), as it possesses the molecular specificity of optical techniques and the penetration depth of ultrasound imaging. However, the precise maneuvering and function control of microrobots, in particular in living organisms, demand the combination of both anatomical and functional imaging methods. Therefore, herein, we report the use of a hybrid High-Frequency Ultrasound (HFUS) and PA imaging system for the real-time tracking of magnetically driven micromotors (single and swarms) in phantoms, ex vivo, and in vivo (in mice bladder and uterus), envisioning their application for targeted drug-delivery.

## 1 Introduction

Micro- and nanorobots (MNRs) offer the potential to operate inside the living body for various healthcare applications,^[1–3]^ as they possess the capacity of reaching hard-to-access locations, non-invasively. Applications such as targeted drug delivery,^[4–7]^ biopsy,^[8]^ blood clot removal,^[9]^ or cell transport^[10]^ have been addressed, for example, by functionalizing the MNRs with biomolecules (e.g. bioreceptors or contrast agents), or by employing smart materials and designs to effectively transport and release a cargo (e.g. cells or drugs).^[1,11–17]^ Some of these micromotors have already been examined in ex vivo and in vivo environments. However, there are still significant limitations when steering single or swarms of MNRs in living organisms,^[2]^ in particular when the intended application and micromotor type require high spatiotemporal resolution with precise anatomical positioning.

Each imaging technique has its own limitations and therefore, researchers are rendering particular efforts to develop or implement imaging systems that could enable precise localization and control within biological tissues. So far, various imaging modalities have been evaluated for the monitoring of MNRs, including ultrasound (US), magnetic resonance imaging (MRI), positron emission tomography-computed tomography (PET-CT), and single-photon emission computed tomography (SPECT), in phantom, ex vivo or in vivo scenarios.^[18–23]^ MRI, for instance, has better imaging contrast for soft tissues than other conventional techniques, but its spatiotemporal resolution is insufficient to visualize small MNRs in real time (below 100 μm). US is suitable for high imaging depth but suffers from a lack of molecular specificity and image contrast. Besides, CT provides deep tissue penetration but lacks temporal resolution and long-term exposures might harm the organism. PET and SPECT provide high sensitivity and molecular information, but the radiation dose remains the foremost concern when an extended use is required to monitor MNRs. Additionally, the optical methods including fluorescence,^[24]^ reflection-based IR imaging^[25]^ or optical coherence tomography (OCT),^[26]^ have been used to track MNRs below scattering tissues with excellent spatiotemporal resolution but have been limited to sub-skin level or superficial medical applications (typically ~1-2 mm in thick biological tissues).^[27,28]^ Unfortunately, in optical methods, spatial resolution degrades significantly with depth due to pronounced light scattering.

To preserve the spatial resolution and molecular specificity of optical techniques while enhancing the temporal resolution and penetration depth, scientists have developed a hybrid optical-ultrasound imaging technique, also called photoacoustic imaging (PAI).^[28–32]^ PAI is based on the photoacoustic effect first observed by Alexander Graham Bell in 1880,^[33]^ which in those days described the generation of sound waves by sunlight. Nowadays, PAI is based on short infrared (IR) laser pulses which are used to excite tissues, inducing thermoelastic expansion/contraction that produces ultrasonic waves.^[34]^ Such ultrasonic waves travel through the tissue until being captured by ultrasensitive ultrasound detectors.^[28,29]^ The use of PAI for the visualization of moving medical microrobots was first suggested by us in 2017.^[2]^ Then we visualized in real-time single magnetically-driven conical micromotors (up to 100 μm long) in three dimensions, underneath ca. 1 cm phantom and in ex vivo chicken tissue samples, highlighting the limits of the technique and showing the advantages of using nanomaterials as labels with unique absorption spectrum to improve micromotors’ image contrast and molecular specificity.^[35–37]^ In later studies, PAI was employed for guiding capsules containing catalytic micromotors in mice intestines, as well as to track swarms of magnetic spiral-like micromotors to treat induced subcutaneous bacterial infection, also in mice.^[23,38]^ Both PAI studies showed the application of micromotors in vivo but there was no clear observation of single micromotor imaging in real-time in living mice, or the distinction between the structure and function of a single or few micromotors and their surrounding environment. Thus, the complementary particulars such as morphology, functionality, and molecular composition are required to enhance the position accuracy of employed MNRs deep in tissue. In this study, we investigate the use of HFUS and PA imaging to monitor swimming micromotors’ motion and function. In particular, we monitor a single or swarm of magnetically-driven spherical micromotors in different scenarios: phantoms, ex vivo, and in vivo (in mouse bladder and uterus). Real-time monitoring of bladder catheterization and the localization of the uterus to monitor micromotors during administration is also demonstrated. Moreover, we perform multispectral imaging to extract the specific absorption spectrum from each of the present entities (MNRs and biological tissues), being of crucial importance when operating in biological which contain a variety of chromophores or absorbing molecules which that hinder the MNRs signal. Finally, we monitor over time the release of a model drug from the spherical micromotors for further drug delivery applications.

## 2 Results and Discussion

### 2.1 Static imaging of single micromotors

A multi-modal HFUS and PA system equipped with a linear array ultrasound transducer and fiber optic bundles on either side of the transducer for illumination (680 to 970 nm) was used for the static imaging of single micromotors (Figure 1a). The micromotors (SiO_2_ particles with a diameter of 100 μm) were fabricated using a drop-casting method followed by the evaporation of subsequent thin metal layers (Ti=10 nm, Fe=50 nm, and Ti=10 nm) (refer to Experimental section). The particles were half-coated with metal layers by electron-beam deposition (Figure 1b-i). The spacing among micromotors can be controlled by varying the concentration of the particles before drop-casting. Figure 1b-ii shows the optical tracking of single/swarm of micromotors when actuated using an external hand-held magnet. The same micromotors can be actuated with rotating magnetic fields as it was previously demonstrated by our group.^[25]^ Figure 1b-iii shows an SEM image of the half-coated particle, highlighting the metal layer surface, and the functionalized particle using a fluorescent anticancer drug, doxorubicin (DOX), for further drug-release monitoring via PAI illustrating the for further in vivo targeted therapeutics (Figure 1b-iv). Initially, we imaged static micromotors to evaluate the performance and ability to locate single of them by HFUS and PA system. The noted number of micromotors was collected using a micro-pipette and then sandwiched between two parafilm sheets. First, bright field (BF) images of the micromotors embedded within the parafilm layers were obtained. Then, PA signal amplitudes of the micromotors were recorded resulting in a good agreement with the previously obtained BF images, with matching locations (Figure 1c-i, ii). The absorbing metal layers provided a broad PA signal in the NIR region (with an excitation wavelength of 800 nm), where the fluence was set below the Maximum Permissible Exposure (MPE) limit (20 mJ/cm^2^) followed by the safe exposure guidelines.^[39]^

**Figure 1.**
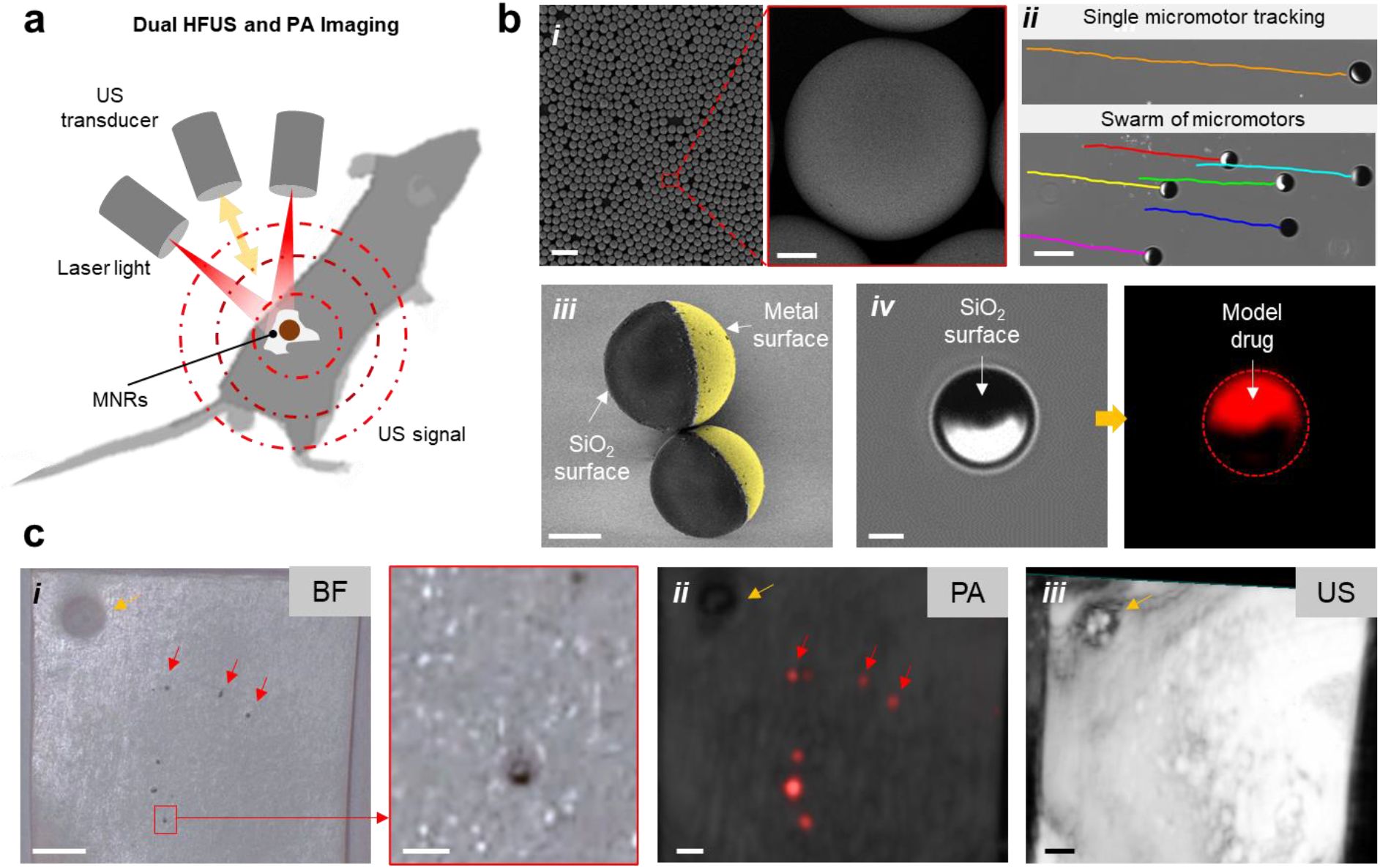
Fabrication and characterization of magnetically-driven spherical micromotors; a) Schematic showing the working principle of the employed hybrid HFUS and PA imaging technique. b) Fabrication and functionalization of spherical micromotors: drop-casted SiO_2_ particles (⌀ = 100 μm) were half-coated with metal layers (10 nm Ti, 50 nm Fe and 10 nm Ti) using electron beam evaporation. Scale bar: 300 μm. Inset scale bar: 20 μm (i), optical tracking of single and swarm of moving micromotors under a magnetic field strength of ca. 60 mT. Scale bar: 200 μm (ii). SEM image of two half-coated spherical micromotors (iii), and bright field and fluorescence images of the functionalized micromotors with a model drug (DOX) (iv). Scale bar for iii, and iv: 40 μm. c) Comparison among BF (i), PA (ii) and HFUS (iii) signal of single micromotors. HFUS shows very strong reflection from the parafilm sheet making micromotors less visible while PA shows the same distribution of individual micromotors in agreement with the BF image data. These images were acquired at 800 nm, where the fluence was set below the MPE limit of 20 mJ/cm^2^. Scale bar: 1 mm.

Figure 1c-iii shows the US image of the embedded micromotors in the parafilm sheet showing no additional information. This phenomenon is explained as follows: HFUS transducers detect high-frequency sound waves (called echoes) to generate images of target objects, and the beam energy provides information about the target along the beam path. The detected echo pattern typically provides information of the tissues or the materials over a selected region of interest. No echoes are generated when sound waves encounter an anechoic structure along the beam path and US images appear black.^[40]^ The intensity of the reflected echoes determines the brightness of the final image. The data from returning echoes are then used to determine the magnitude of each echo received, and the distance of the target object from the transducer position. Strongly attenuating objects like bones reflect echoes and cause a dark tail like shadow.^[40]^ However, strong scattering of echoes from the two-layer parafilm sheet decreases significantly the image contrast and masks the micromotor signal, making the parafilm platform not suitable for US imaging experiments.

### 2.2 Tracking of single micromotors in phantom and ex vivo tissue

To minimize the artifacts from HFUS scattering due to the employed parafilm-based sample holder, the following experiments were conducted in both tubing phantom and ex vivo tissues. For the tubing phantom setup, a commercially available methacrylate support (Vevo Phantom, FUJIFILM VisualSonics, The Netherlands) was used to mount transparent intravascular polyurethane (IPU) tubing (inner diameter ~380 μm and outer diameter ~840 μm, SAI Infusion Technologies, USA) as a fluidic channel for flow experiments. The micromotors were inserted in the tube, and it was then immersed in the phantom chamber containing DI water for better acoustic coupling (Figure 2a). After the PA image acquisition, the spectral characteristics of the samples were recorded. 3D single-wavelength (800 nm) imaging was also performed using a 2D stepper-motor and a linear translation of the transducer over the IPU tubing. For control experiments, another tube was mounted with a saline solution. All PA measurements were performed at a gain of 40 dB.

**Figure 2.**
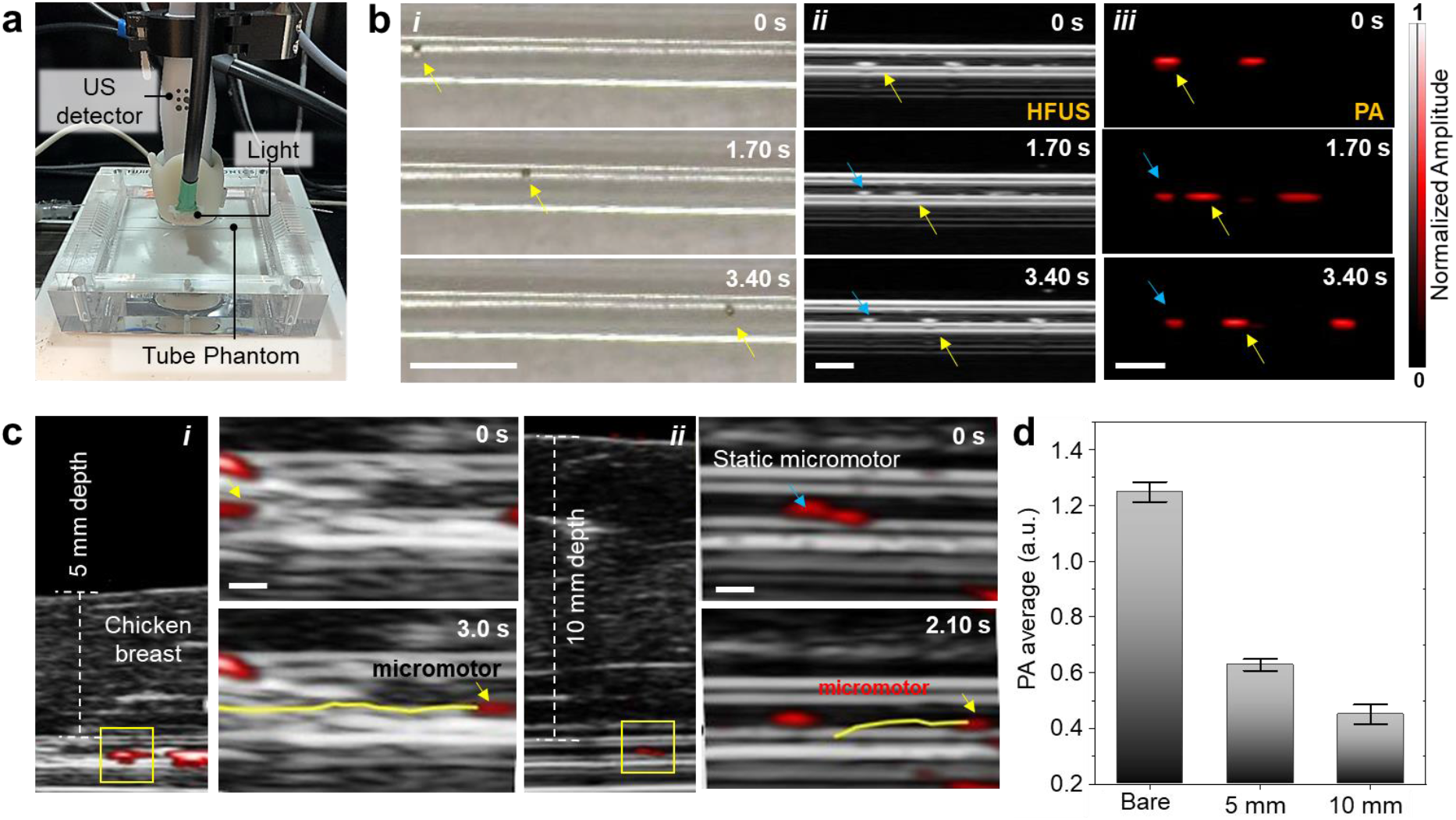
Tracking of single micromotors in phantom and ex vivo chicken tissues; a) Phantom chamber with tubing immersed in a water batch with HFUS-PA detector on top of it for real-time tracking of moving micromotors. b) Time-lapse images of a 100 μm moving micromotor in BF (i), HFUS (ii) and PA (iii) modes. Yellow arrows show the single moving micromotors. Scale bars: 1 mm. c) Micromotors embedded within chicken breast tissue and the time-lapse images of a moving micromotor below 5 mm (i) and 10 mm (ii) thick tissue. Scale bar: 500 μm. d) PA signal amplitude of single micromotors with comparable ROIs below 5- and 10-mm thick tissue, which decreases with increased penetration depth. Blue arrows point at static microstructures and are used as reference points.

US image quality is influenced by the medium through which sound travels from the transducer. Material choice is a crucial factor in US imaging to match the speed of sound and to avoid undesired artifacts. Thus, the materials must possess low attenuation, meaning that acoustic waves are able to reach the target (micromotor) and return. The material should also exhibit low backscatter between the micromotor and the transducer. The ratio of the micromotor reflection intensity to backscatter intensity is chosen as a relative figure-of-merit to determine the material performance.^[41]^ On the other hand, parafilm, as a semi-transparent sheet, exhibited high front surface reflections which reduced the signal intensity from the embedded micromotors to the transducer. While the IPU tubing features low attenuation and low front surface backscatter that provides a better HFUS contrast making possible to visualize single moving micromotors inside this tube phantom. The dynamic tracking of a single micromotor is shown in Video S1 and in time-lapse images from 0 to 3.40 s (Figure 2b), comparing BF, HFUS, and PA imaging modes. Blue arrows show the position of static micromotors (stuck on the tube walls) while the yellow arrows correspond to the single moving micromotor. The overlaid HFUS-PA video shows single micromotor moving forward and backwards with clear localization within the phantom tube (Video S2). This hybrid feature is useful in deep tissue imaging where US provides tissue background morphology and PA maintains high contrast of the micromotors. The measurement was performed with a position-fixed high frequency transducer (21 MHz) to avoid image distortion.

To visualize the micromotors underneath biological tissues, we employed chicken breast from a local store, with an approx. 5- and 10-mm thickness (measured with a Vernier caliper). The IPU phantom tubing containing micromotors was placed in between two layers of chicken breast tissue (Figure S2) for the consequent measurements. The US gel was applied between the detector surface and the top of the tissue to match the acoustic impedance for efficient signal transfer. HFUS brightness-mode (B-mode) generated a structural image, suitable to identify the tubing structure, and the PA system was used to monitor the displacement of the micromotors, by detecting their light absorption characteristics. Both images were acquired in real-time during external magnetic steering. The locomotion of a single micromotor was tracked under 5- and 10 mm thick tissue samples and it was possible to display overlaid HFUS and PA images as shown in time-lapse images (Figure 2c and Video S3). The corresponding PA amplitude values of the micromotors are shown in Figure 2d. The micromotors showed decreased PA intensity when increasing the tissue thickness due to pronounced scattering of the excitation light in deep tissues.

### 2.3 Micromotors swimming in mouse bladder and uterus

The following in vivo HFUS and PA experiment was performed under the animal handling license No. DVS06, hold by the co-authors of FUJIFILM VisualSonics, The Netherlands, ensuring the biocompatibility nature of the employed materials. The in vivo study of the micromotors was conducted using the same equipment and the imaging settings as those for in phantoms or ex vivo studies. The animal testing was performed to illustrate the potential of hybrid HFUS and PA imaging system regarding penetration depth and real-time navigation ability of a single or swarm of micromotors. The micromotors were injected into the bladder and uterus of 12-week old mice (details in the Experimental Section). We selected the bladder and uterus cavity as sites of injection to allow enough room for the micromotors free swimming. The bladder is a hollow soft organ that serves as a reservoir for the storage and periodic release of urine, with areal dimensions of approx. 5.50×6.0 mm as shown in 2D US image with anatomical features of bladder (Figure S3). Under anesthesia, the mouse bladder was catheterized by following standard safety guidelines and protocols.^[42]^ Before starting the experiments, the physiological parameters of the mouse were monitored (Figure S4 and Video S4). First, the micromotors were collected (15-25 in ~20 μl PBS) in a catheter, and then inserted into the bladder. The catheter was inserted through the urethra of an anesthetized mouse as shown in schematic in Figure 3a. The position of the catheter and bladder was monitored in vivo as shown in the 3D B-mode reconstructed image (Figure 3b). After reaching the entrance of the bladder, the micromotors were released gradually into the bladder cavity. The overlaid HFUS and PA image in Figure 3c, shows the micromotors near the catheter edge before and after release. PA signal (red spot) highlights the swarm of micromotors being ejected from the catheter. While US image highlights the anatomy of the organ of operation, the PA image shows the molecular information (meaning the material absorption properties) of the imaged micromotors. After release, the micromotors started to swim down freely in the urinal fluid, and it was possible to visualize the micromotors from a single to very few and even a swarm of them while sinking to the bottom of the bladder cavity (Video S5). Time-lapse HFUS and PA images show the swimming behavior of micromotors in urinal fluid from top to bottom surface of the bladder over a period of 2.50 s (Figure 3d) and the swimming speed or falling velocity of a cluster of micromotors in the bladder was estimated to be ca. 1250 μm/s (in the current experiment). The downward displacement of one of a small cluster was ca. 1.6 mm which took 1.3 s to reach the bottom of the bladder. Finally, the whole swarm of micromotors at the bottom of the bladder was manipulated by using the external magnet and the swimming micromotors were visualized within the bladder environment (Video S6). The upward displacement of the whole swarm centroid was estimated to be ~3.5 mm (in the current experiment) after a time interval of ca. 8 s and the dimension of the moving cluster was estimated to be between 2 to 3 mm wide (Figure 3e). The upward speed of the swarm was found to be ~480 μm/s after time travel of 7 s and as expected, the speed of the cluster in opposite direction (upward motion) was slower than the downward motion as the gravitational force was dominant over the drag force (with no applied magnetic field). The speed of single or swarm of micromotors also depends on the strength of the applied field at the evaluated distance from the micromotor position. In this experiment, the magnet was place directly onto mouse skin, after removing the rodent fur with a depilation cream, being approx. 2 cm away from the micromotors location (inside the mouse bladder), resulting in an estimated magnetic field of ~60 mT. (Figure S1). It is worth noting that this is a proof-of-technology study that shows the ability of controlled movement of micromotors in real-time in mice by employing HFUS and PA imaging. However, a more sophisticated setup including dedicated magnetic field actuators are necessary for a precise steering and complex actuation of the here-presented micromotors and can be extended for any other medical micromotor.

**Figure 3.**
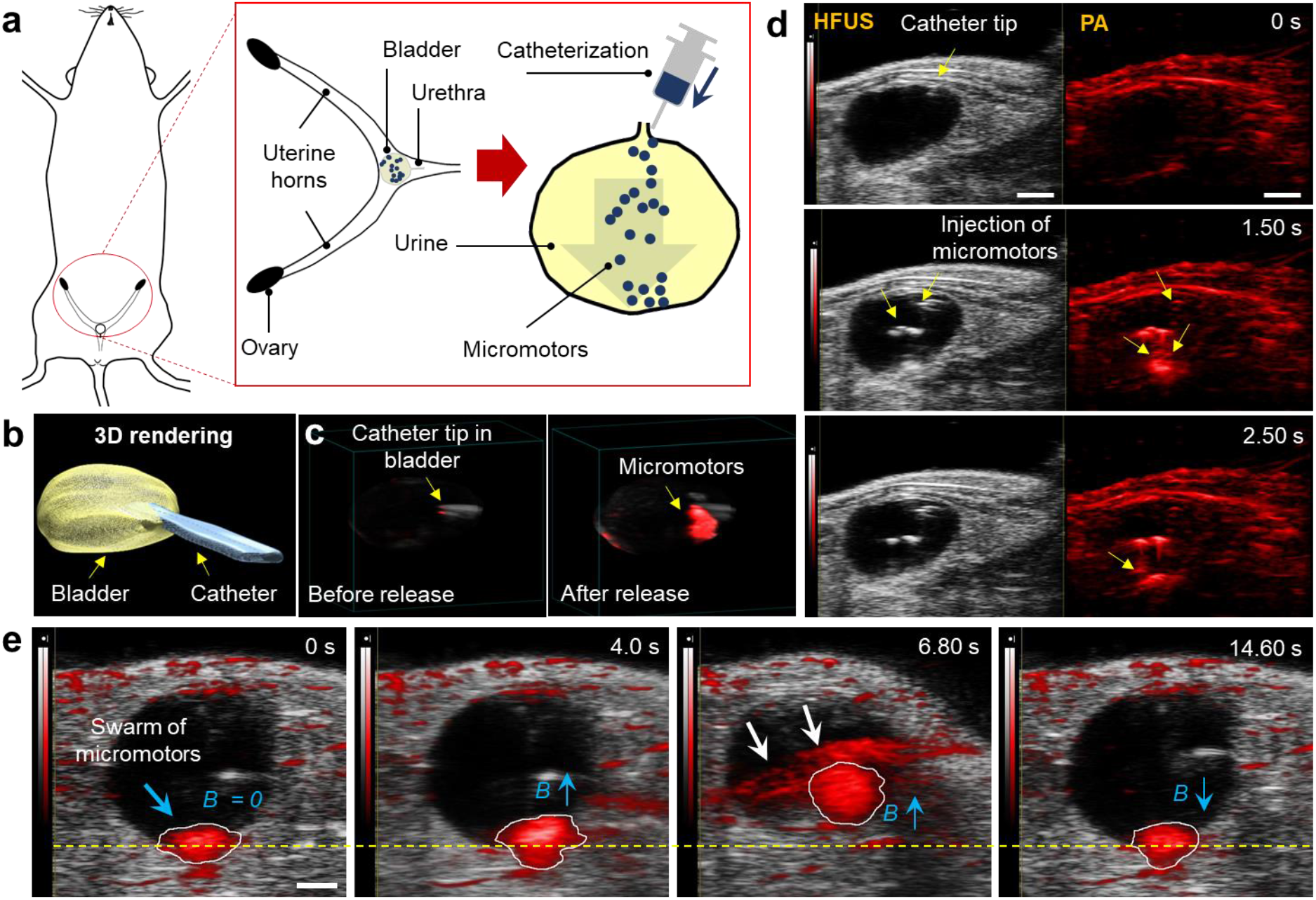
HFUS and PA imaging of a single and swarm of micromotors swimming inside a mouse bladder; a) Schematic showing the position of the bladder. b) 3D reconstruction of the bladder with the inserted catheter, after performing the catherization via real-time US feedback. c) HFUS and PA overlaid image showing the location of the catheter tip, before and after releasing the micromotors. d) Hybrid HFUS and PA time-lapse images of swimming and free-falling micromotors over a time period of 2.50 s. The micromotors migrate downward after being released from the catheter tip individually and then accumulate at the bottom surface. Scale bar, 1 mm. e) A swarm of swimming micromotors under the actuation of an external hand-held magnet (estimated magnetic field strength (~60 mT)). The upward displacement of the swarm centroid from the dashed yellow reference line, was estimated to be ~3.5 mm after a time interval of approx. 8 s. The dimension of this moving cluster was ca. 2 to 3 mm wide. White arrows show the artifacts arising from the close proximity of the external hand-held magnet. Scale bar, 2 mm.

Apart from bladder, we also evaluated the micromotors imaging in the mouse uterus. Here, the catheter was inserted via the vagina and cervix into the uterus cavity by following the previous safety guidelines.^[42]^ In this experiment, we investigated the smallest detectable feature size of micromotors in uterus by injecting controlled number of micromotors as depicted in the schematic in Figure 4a. The samples were dispersed in PBS containing few micromotors (<10) in a catheter. By employing US imaging, it was possible to precisely locate the catheter tip at the opening of the uterus cavity prior micromotors insertion (Figure 4b). Ultimately, it was possible to deliver ca. 2-3 micromotors in the mouse uterus from the catheter needle. One of the micromotors was steered in real-time in the uterus channel over a trajectory of ~950 μm (Video S7). Time-lapse HFUS and PA images showed the moving micromotors from one position to another over a period of 3.60 s (Figure 4c). The resulted images comprise PA amplitude values of the uterus cavity with and without micromotors (control) by choosing comparable ROIs (Figure 4d).

**Figure 4.**
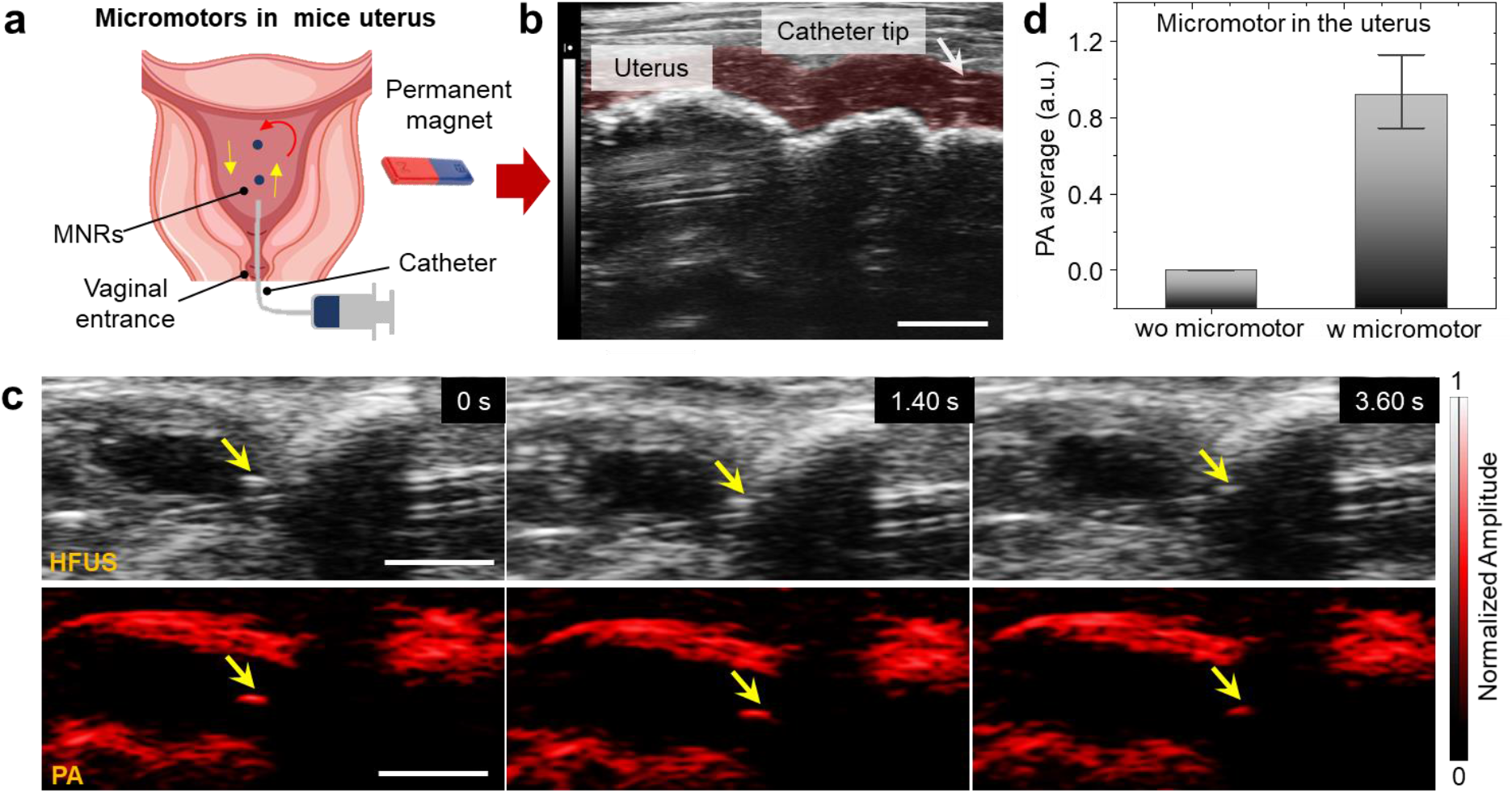
HFUS and PA imaging of micromotors moving in the uterus; a) Schematic showing the insertion of the catheter into uterus body through the vagina and moving micromotors by applying an external magnetic field. b) Real-time HFUS feedback positioning of the catheter in the mouse’s uterus. Red-shaded area indicates the uterus channel. Scale bar is 2 mm. c) Time-lapse images of single moving micromotors (Yellow arrows show the position of moving micromotor). Scale bar is 2 mm. d) PA amplitude values extracted from the captured PA images of a single moving micromotor, compared to the control (uterus location where no micromotors was visible “wo micromotor”).

HFUS images have speckle patterns from the background tissues which make it challenging to identify the micromotors. The tracked micromotor was visible in both HFUS and PA modes, however it is clear that HFUS provided a better visualization of the catheter entrance and the surrounding uterus anatomic features, while PA facilitated the discrimination of the light absorbing micromotors from the surrounding environment, offering a high SNR ratio. The employed micromotors and the here-evaluated hybrid imaging system is appealing for the supervised drug cargo-delivery towards urinary tract diseases e.g. urethral stenosis, bladder cancer, or infection, or towards in vivo assisted fertilization, where similar engineered parts can be used to guide or transport sperm.^[4,10]^

### 2.4 3D multiplexing in vivo

The advantage of multiplexed imaging or spectral unmixing is in being able to distinguish light absorbing signals coming from the different organic and inorganic components present in the field of view.^[43]^ The multi-wavelength mode takes advantage of the tunability of the pulsed laser source to acquire PA data at multiple wavelengths in 2D and 3D for automated post-processing, obtaining a unique spectrum. Such specific spectral capacity of PAI offers specific detection of endogenous molecules like oxygenated (oxy-Hb) and deoxygenated hemoglobin (deoxy-Hb), among others.^[29]^ The first step is to image the target (micromotors, oxy- and deoxy-Hb or other employed materials) at multiple wavelengths. Then spectral unmixing algorithm is applied, generating separate images based on the contrast obtained from the different absorber materials (endogenous or exogeneous). Thus, when merging the independent images, one can obtain the distribution of oxy- or deoxy-hemoglobin as well as of the present micromotors marked according their absorption properties.

In this experiment, the mouse was injected with a 10 μL mixture of PBS and target micromotors into the hind limb (popliteal lymph node fat pad), followed by the PA multi-wavelength measurement in the wavelength range of 680 to 970 nm with a step size of 5 nm in the spectral mode, at a frequency of 30 MHz. The obtained signals were then spectrally un-mixed to discriminate the PA signal of injected micromotors from the surrounding tissue chromophores (Figure 5a). 3D reconstruction of the target site demonstrates the spectral behavior of the injected micromotors (yellow color) and the chromophores present in blood i.e. oxy-Hb, deoxy-Hb (Figure 5b). For deep in vivo imaging, it is important that the introduced micromotors hold a unique absorption signature compared to the tissue background. Figure 5c shows the PA amplitude of the micromotors (yellow curve), in comparison with oxy- and deoxy-Hb and it is possible to clearly distinguish the injected micromotors. The PA amplitude values of the micromotors were extracted from the captured PA images and then calibrated with measured optical absorption of hemoglobin. 3D multiplexing PA scans have allowed investigating the biodistribution of the injected micromotors in vivo which in turn can be used to target specific body organs to perform a certain medical task.

**Figure 5.**
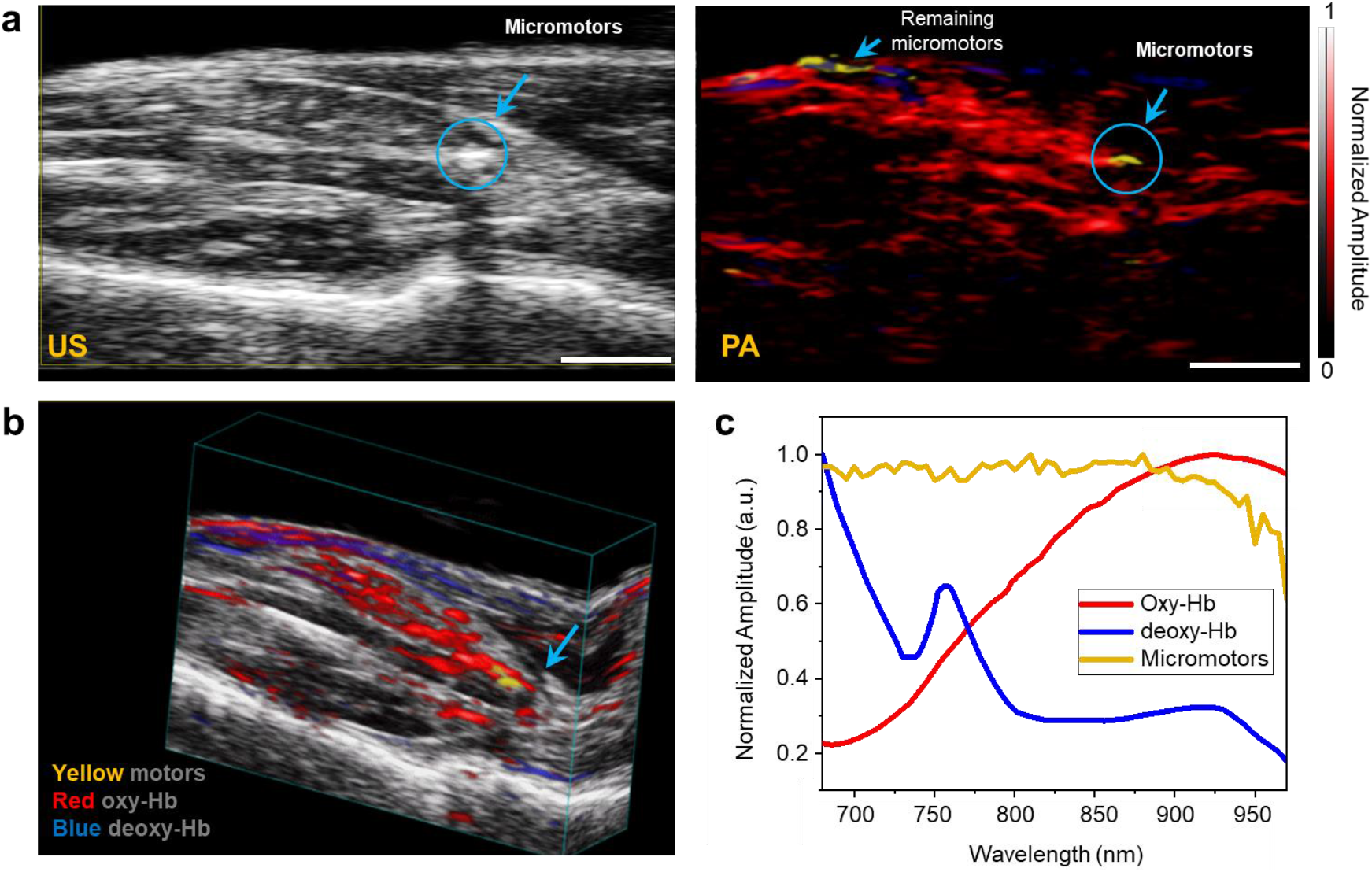
3D multiplexing in vivo; a) Injection of micromotors into the mouse hind limb (popliteal lymph node fat pad) and HFUS and PA imaging of a swarm of micromotors (Yellow spot in PA mode shows the position of injected micromotors within tissue). The remaining micromotors from syringe tip were also visible on tissue top surface after taking it out. b) 3D reconstruction and spectral unmixing of the injected micromotors in tissue. c) PA signal strength of the injected micromotors (yellow), oxy (red) and deoxygenated (blue) hemoglobin.

We used metal-coated micromotors that absorb light and generate spectral signatures for acoustic detection and the resulting PA signals exhibited a broad absorption spectrum. In the uterus and bladder cavity, due to the fluid inside, image contrast was better to distinguish the micromotors. However, when injected within the tissue, the signal from oxy- and deoxy Hb, and other chromophores hindered the signal of the micromotor. In this case, spectral unmixing was applied to distinguish the signals by their specific absorption peaks. Another strategy to enhance the PA signal of injected micromotors is to stamp them with suitable and biocompatible contrast agents that absorb in a narrow and specific wavelength range, such as the ones shown previously by our group^[35,36]^ with a strong absorption peak in the near infrared range.

### 2.5 Towards targeted drug delivery monitoring

Here, we propose a potential scenario where this technique and the employed spherical micromotors could be applied. As reported previously in the literature, metal-coated particles have been shown as drug carriers, propelled either by bacteria or by chemical reactions to target cancer cells in vitro.^[25,44,45]^ The loading mechanism ranges from physical absorption to more sophisticated stimuli-responsive delivery triggers. To evaluate the feasibility of HFUS and PA to not only identify the drug-loaded micromotors but also to monitor the drug release overtime, we functionalized the micromotors with a model biomolecule (indocyanine, ICG), similar as it was done for DOX (Figure 1b-iv). ICG, an FDA approved contrast agent, was chosen due to its unique absorption peak at 880 nm which facilitated their visualization via PAI, with negligible artifacts arising from autofluorescence or surrounding tissues deep in the body. As a proof of concept, we first functionalized the spherical micromotors by overnight incubation in an ICG solution (details in the Experimental Section). The illustration of ICG or a drug immobilized onto the SiO_2_ surface of the Janus micromotor, followed by the drug release mechanism over time is shown in Figure 6a. To test if the cargo was successfully loaded onto the micromotor, a solution of ~20 μl of ICG-loaded micromotors was injected into IPU tubing and HFUS and PA signals were recorded in static conditions, as shown in Figure 6b. An ROI was chosen to plot the PA spectrum of ICG-loaded micromotors. ICG-loaded micromotors exhibited a distinct absorption peak at around 880 nm as expected from its optical properties (Figure 6c, red curve). After a time-lapse of ca. 2 min, the same ROI was analyzed to get the PA signal of ICG-loaded micromotors, showing a decrease in PA amplitude without any distinct peak, indicating the complete release of ICG, which rapidly diffused in the surrounding after NIR light was turn on (Figure 6c, blue curve). This test highlights the advantages of the PA system to monitor known optical signatures of implemented materials/drugs for precise cargo-delivery inspection. Different drug release mechanisms can also be explored using this technique, as in a recently reported work, where the NIR light irradiation by PAI was sufficient to disintegrate the capsules to deliver the micromotors in the stomach of a mouse,^[38]^ or for triggering the drug release for treating sub-skin pathogenic bacterial infections via photothermal therapy (heating of a bacterial site up to 54 °C).^[23]^

**Figure 6.**
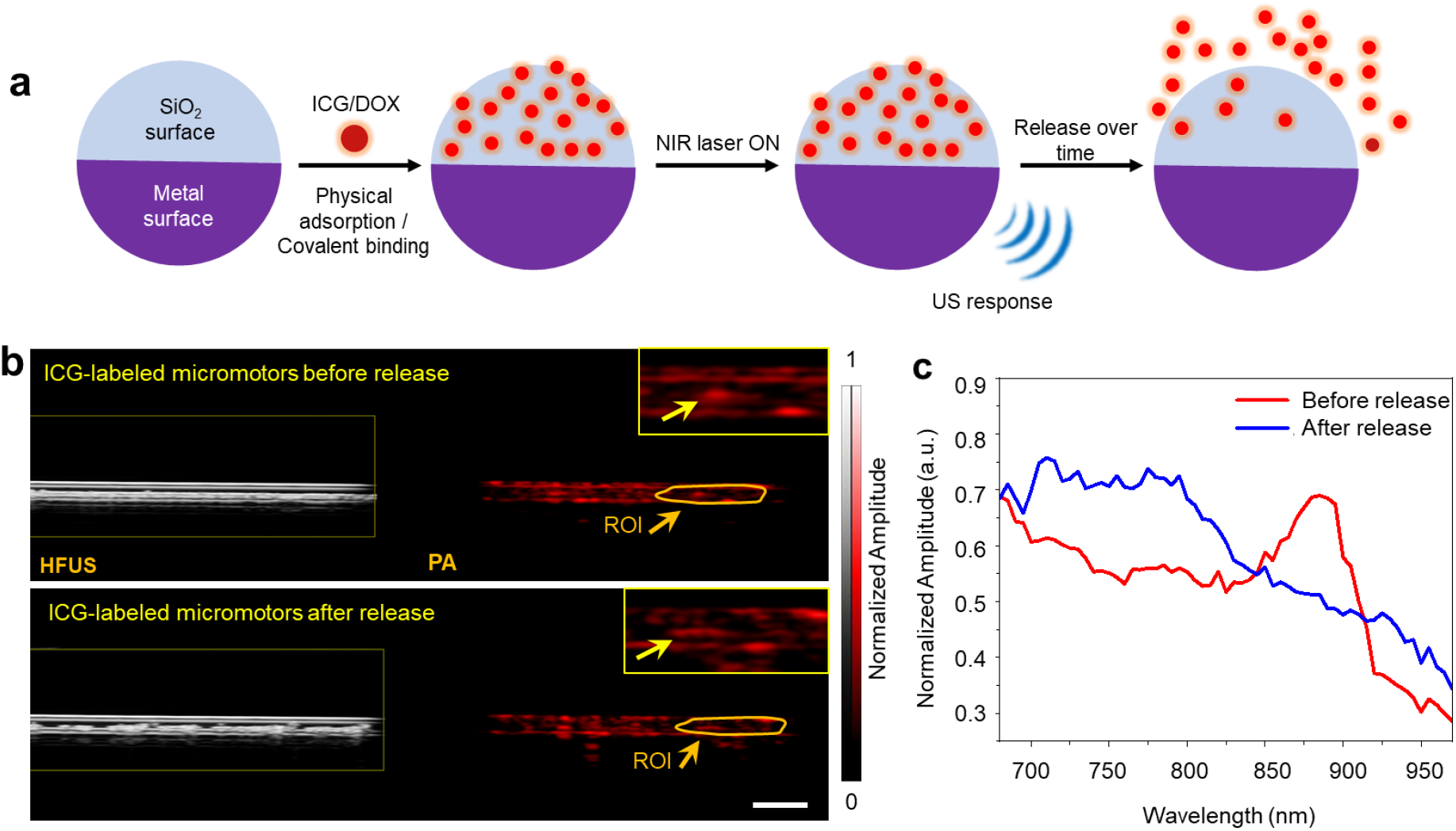
Application scenario of micromotors for drug delivery monitoring; a) Schematic showing the immobilization of a model drug on the Janus micromotor, followed by its release. b) ICG-coated micromotors injected into IPU tubing. Scale bar is 1 mm. c) PA spectral signal showing a distinct absorption peak at around 880 nm (red curve) before and after release, when analyzing a fix ROI where a few static micromotors were located.

Future work will be focused on more complex positioning of the medical micromotors in the living mice by external magnetic fields and on the control and monitoring of the cargo-release both by the employed HFUS and PA technique.

## 3 Conclusions

Hybrid imaging systems, in particular when combining anatomical and functional operating modes provide superior advantages for translational clinical and biomedical studies. For example, dual PET-CT system offers a combination of high-resolution images as well as intravital molecular data.^[46]^ Similarly, hybrid fluorescence molecular tomography and X-ray CT offers more diversity than operating them alone e.g. improved penetration depth.^[47]^ In the present study, we implemented a hybrid HFUS and PA imaging system to carry out functional and structural imaging of swimming micromotors deep in the bladder and uterus of a mice model. Single or few micromotors were successfully visualized in phantoms, ex vivo, and in vivo environments. US imaging showed clear boundaries among different types of organs and precisely identified the position of the catheter (with micromotors) in the body with real-time feedback, improving the accuracy of the procedure and avoiding tissue damaging during catherization. In US images, internal structures like a bladder and uterus were visible to inject the micromotors while the PA images displayed the optical absorption characteristics of tissues and microstructures in mice.

Our results showed that the hybrid imaging mode provided from one side the volumetric US image and the PA distribution map of hemoglobins in the region of interest and resolved the micromotors from the signal of the surrounding tissues. The spatial resolution conferred by the HFUS and PA probe facilitated the visualization of micromotors. HFUS was suitable for the real-time monitoring of the catheterization and localization of the target organ to deliver the micromotors, while PA allowed a reliable monitoring of the injected micromotors. The sensitivity of the hybrid system in the tissue enabled it to capture and discriminate the distinct optical contrast between endogenous chromophores and the micromotors, being of crucial importance when identifying MNRs position in a complex biological environment with poor contrast.

So far, the here reported experiments have been performed in ex-vivo tissues and in living mice. However, to translate this technology to humans and despite all the ethical discussions around the employment of medical microrobots which will arise in the next years, there are some technical limitations which need to be solved such as the penetration depth (so far ca. 2 cm, with micrometric resolution has been possible), but going beyond this limit will compromise the spatial and temporal resolution. Biodegradability is another prerequisite. There have been few attempts of biocompatible micromotors for in vivo drug-delivery applications^[38,48]^ and eventually, the micromotors should clear the body after performing an assigned task without any diverse effects. A preliminary study of micromotors using hybrid HFUS and PA system and an envisioned application in the field of supervised cargo-delivery is presented.

In summary, the hybrid HFUS and PA system is a powerful tool in addressing various imaging challenges of micromotors that require correlation with molecular and biological data, opening up a new horizon for imaging of medical microrobots in living organisms.

## 4 Experimental Section

### 4.1 Fabrication of the micromotors

The micromotors were constructed using a drop casting method with subsequent thin metal layer deposition. First, glass substrates (22×22 mm²) were rinsed and ultra-sonicated (Elmasonic, Elma Schmidbauer GmbH) in acetone and in isopropanol for 3 min each and finally dried in a stream of N_2_. The substrates were exposed with oxygen plasma (Diener electronics), leading to the removal of impurities and contaminants from glass surface, to obtain clean and hydrophilic substrates. Afterwards, a monolayer of micro-sized (⌀ = 100 μm, Corpuscular Inc. USA) silicon dioxide (SiO_2_) particles was assembled. In detail, SiO_2_ particles were washed with methanol and centrifuged for 1 min to remove supernatant and then again mixed with methanol and vortexed (Vortex mixer, VWR) prior usage. Silica particles were mixed thoroughly in methanol and ~20-25 μl of particle-solvent solution was drop-casted on the edge of a pre-plasma treated glass slide. The glass slide was adjusted at an angle to allow the spreading of silica particles over the entire glass cover slip, making a homogeneous monolayer of silica particles. The micro arrays were randomly formed in the direction of solvent evaporation. The resulted monolayer was dried at room temperature. After drying, the samples were half-coated with Ti (10 nm), Fe (50 nm) and Ti (10 nm) of high purity (99.995%) by electron beam physical vapor deposition (e-beam evaporator, Plassys MEB550S) at a deposition rate between 0.5-1.0 Å/s. Fe layer was evaporated for magnetic guidance and Ti was chosen for biocompatibility.

SEM and optical microscopy were performed for the characterization of the prepared micromotors. Scanning electron microscopy (SEM, Zeiss Nvision 40, Carl Zeiss Microscopy GmbH) was performed by coating the sample with ~10 nm Pt to make the specimen conductive and to avoid charging effects during imaging. Bright-field microscopy was performed using an optical microscope (Zeiss Microscopy GmbH) equipped with 5x, 10x and 20x objective. Origin 2019 and ImageJ software were used for the analysis of the experimental data.

### 4.2 Ex vivo tissue set-up

Chicken breast samples were purchased from a local store and used with thicknesses of 5 and 10 mm. The IPU tubing was filled with the micromotors suspended in PBS 1X and then placed between two 5 mm tissue samples. The process was repeated for 10 mm thick tissues afterwards. To ensure uniform distribution of thickness and prevent from drying, the thickness of the tissue samples was measured using a digital Vernier Caliper. The tissues containing micromotors were then placed on an imaging chamber grid for tracking measurements.

### 4.3 In vivo mice handling

The in vivo mice experiments were conducted at the FUJIFILM VisualSonics laboratory in Amsterdam. The animal protocols used in this work were appraised and approved by the Committee on Ethics in the Use of Animals (CEUA), The Netherlands (Protocol AVD2450020173644). They are in conformity with FELASA guidelines and the National Law for Laboratory Animal Experimentation (Law No. 18.611). The experiments were performed by using the same equipment used for the phantom studies. 12 weeks old mice were used for in vivo imaging in bladder and uterus. All mice were anaesthetized using isoflurane, at a concentration of 1.5-2%. O2 flow (1>2 mL/min maintenance) mixed with isoflurane. For optimized coupling of the ultrasound gel the rodent fur needed to be removed using commercially available depilation cream. For 3D multiplexing imaging, an ICR(CD-1) +/− 25-gram female mouse was injected with a 10 μL mix of PBS 1X and micromotors into the hind limb (popliteal lymph node fat pad). 3D PA multiwavelength was applied to automatically un-mix the injected micromotors. PA multispectral imaging distinguished the difference in optical signals related to the micromotors in 2D and 3D for subsequent post-processing.

### 4.4 HFUS and PA imaging of the micromotors

PA measurements were carried out by using the Vevo-LAZR X (FUJIFILM VisualSonics, The Netherlands) system, a multimodal platform which allows the simultaneous imaging of high-resolution ultrasound and photoacoustics. The system was equipped with a linear array ultrasound transducer at a central frequency 21 MHz (MX 250, FUJIFILM VisualSonics, The Netherlands) and fiber optic bundles on either side of the transducer for illumination. The fiber bundle was coupled to a tunable Nd: YAG laser (680 to 970 nm) with a 20 Hz repetition rate and the signals were collected by the 256-element linear array transducer. The pulsed laser generated a wavelength-tunable pulsed beam (680-970 nm) which was delivered by a bifurcated fiber bundle integrated with US transducer. Both HFUS and PA signals were collected and reconstructed using on-board software. For single pulse excitation, the PA images were acquired at an excitation wavelength of 800 nm (with in-plane axial (75 μm) and temporal (5-20 fps) resolution. The same protocol was applicable for in vivo imaging in mice bladder and uterus.

### 4.5 Drug loading (ICG or DOX)

The spherical micromotors were loaded with an anticancer drug (doxorubicin, DOX), which is widely approved for cancer therapy and a model drug (indocyanine, ICG). ICG is an FDA approved contrast agent for use in humans and was employed for this experiment due to its unique absorption peak at 880 nm. The micromotors were functionalized by overnight incubation in an ICG or DOX using the same protocol. ICG or DOX solution was prepared with a concentration of 100 μg/ml and then mixed with the micromotor solution with PBS (~10 μL). The mixture was covered with aluminum foil and incubated for 24 h.

To evaluate the loading of the DOX micromotors, fluorescence microscopy was used to track a single micromotor with an excitation wavelength of 470 nm). It was possible to distinguish drug-loaded micromotors from un-loaded (control) ones using fluorescence microscopy. DOX possesses an intrinsic fluorescence with an emission peak at around 600 nm (Figure 1b-iv). It shows the potential of using micromotors as a drug-carriers.

## Acknowledgements

This work was supported by the German Research Foundation SPP 1726 “Microswimmers-From Single Particle Motion to Collective Behavior” and for the European Research Council (ERC) under the European Union’s Horizon 2020 research and innovation program (grant agreement No. 835268 and No. 853609). O. G. Schmidt acknowledges financial support by the Leibniz Program of the German Research Foundation. Authors also thank Jithin Jose for his valuable comments and discussion in manuscript writing.

## Notes

The authors declare no competing financial interest.

Received: ((will be filled in by the editorial staff))

Revised: ((will be filled in by the editorial staff))

Published online: ((will be filled in by the editorial staff))

